# Intraspecific diversity based on low temperature induced arrhythmia in embryonic heart of medaka (*Oryzias latipes*)

**DOI:** 10.1101/2022.04.05.487105

**Authors:** Wataru Yoshimura, Shoji Oda, Takafumi Katsumura, Hiroshi Mitani, Tomomi Watanabe-Asaka

## Abstract

**Background:** Cold tolerance during embryonic development, especially blood circulation is important for growth in poikilothermic animals. Medaka (*Oryzias latipes*), has cold tolerance and is distributed in the highest latitudes occupied by the genus *Oryzias*. Regarding cold tolerance in embryogenesis, Hd-rR strain belonging to the southern Japanese (S.JPN) group showed arrhythmia when the embryo was exposed to 15 °C in the heartbeat initiation period (st. 24), whereas the embryo of Odate medaka, which belongs to the northern Japanese (N.JPN) group, showed stable heartbeats.

**Results:** In this study, to verify the sensitivity to low temperature in the medaka wild populations, heartbeat intervals of st. 24 embryos derived from three N.JPN, eight S.JPN, and two western Korean/Chinese (W.KOR) populations were investigated at an arbitrary temperature range of 12–15 °C. There was no significant difference in the mean values of the heart rates and the coefficient of variation (CV) of interbeat intervals in the N.JPN, S.JPN, and W.KOR groups. A temperature dependency of the CV within 12–15 °C was observed only in five S.JPN specimens, whereas three S.JPN, the N.JPN and the W.KOR specimens showed no alteration in CV with temperature. Temperature dependency of the heart rate was also varied in the S.JPN specimens.

**Conclusions:** These results suggest that the temperature dependency of the CV under 12–15 °C during the heartbeat initiation period is a variation within the S.JPN after the divergence from the N.JPN.

## Background

Medaka (*Oryzias latipes*), or Japanese rice fish, is a small freshwater fish distributed in the highest latitudes (Japan, Korea, and China) occupied by the *Oryzias* genera, which inhabit wide areas of East Asia from the tropical to the temperate zone, and tolerate cold climate (Iwamatsu, 1989; Iwamatsu, 2006). The wild population of medaka consists of four genetically and geographically divergent groups: the northern Japanese (N.JPN), the southern Japanese (S.JPN), the eastern Korean (E.KOR), and the western Korean/Chinese (W.KOR) groups (Sakaizumi, 1986; Sakaizumi, 1983; Takehana, 2003; Katsumura, 2009; Spivakov, 2014). A common ancestor of the medaka is estimated to have diverged from a sister lineage *O. luzonensis*, followed by a divergence between the continental (E.KOR and W.KOR) and Japanese populations 5.4–6.0 million years (Myr) ago (Takehena, 2003; Kasahara, 2007). The two Japanese populations (N.JPN and S.JPN) diverged 4–5 Myr ago (Watanabe, 2006; Kasahara, 2007; Katsumura, 2017). These medaka groups have over 80 wild populations with large genetic diversity and have been used for analyses of the functional differences of genetic polymorphisms (Matsumoto, 2009; Katsumura, 2014; Igarashi, 2017).

The heart constantly circulates blood to the body by rhythmic beats. The heartbeat is generated by the sinoatrial node in the right atrium of the heart (Wilders, 1993; Guevara, 1995). External temperature is one of the important factors that affect the heartbeat rhythms (Murayama, 2017). Adult medaka overwinter at 4 °C of minimum water temperature and lay eggs in early spring at 10 °C of minimum water temperature (Shirai, 1937). During embryogenesis at low temperatures, several phenotypes have been reported to suffer developmental arrest at the blastula stage, bradyarrhythmia at heartbeat initiation stage, and regurgitation or clogging of the blood flow at later stages (Shirai, 1937; Iwamatsu, 2006; Watanabe-Asaka, 2014). In medaka embryos, the stage of heartbeat initiation period (st. 24, Iwamatsu 2006) is highly sensitive to low temperature. In a previous study (Watanabe-Asaka, 2014), Hd-rR strain embryos from the S.JPN group showed arrhythmias at st. 24 and blood regurgitation at the heart development stage (st. 36) at 15 °C, whereas Odate medaka embryos from the N.JPN group showed rhythmical heartbeat under similar conditions with normal blood flow in the later stage. It was suggested from a phylogenetic analysis (Takehana, 2003) that the N.JPN group acquired stable heartbeat during embryogenesis at low temperature and expanded in the north because the acquisition of broad temperature adaptability enabled expansion into new environments.

In this study, we further examined the heartbeat stability of st. 24 embryos and temperature dependency in the range of 12–15 °C using the wild medaka lab stocks of three, eight, and two populations derived from the N.JPN, S.JPN, and W.KOR groups, respectively. The coefficient of variation (CV) of interbeat intervals with temperature was categorized in two groups: a “negative-correlation group” of five S.JPN populations and a “less-correlation group,” which includes all the N.JPN and W.KOR groups and three S.JPN populations. These results suggest that the sensitivities to low temperature of medaka embryos at the heartbeat initiation stage differ within populations of the S.JPN, while the N.JPN and W.KOR groups show fewer differences.

## Materials and Methods

### Ethics

All experiments were performed in accordance with Japanese laws and the guidelines for the care of experimental animals according to the University of Tokyo Animal Experiment Enforcement Rules. The protocol was approved by the Committee on the Ethics of Animal Experiments of the University of Tokyo (Permit Number: C-14-02). All efforts were made to minimize suffering.

### Fish maintenance and husbandry

Fish of the inbred HNI strain derived from the N.JPN group were obtained from the National BioResource Project (NBRP) medaka. Fish from the other inbred Hd-rR strain derived from the S.JPN group were obtained from our breeding colony. We also used 13 wild lab stocks derived from local populations: Odate, Kaga, and Maizuru medaka from the N.JPN group; Ichinoseki, Urizura, Okaya, Iida, Tanabe, Arita, Kazusa, and Gushikami medaka from the S.JPN group; and Maegok and Shanghai medaka from the W.KOR group. Wild lab stocks from local populations used in this study were provided by the Graduate School of Frontier Sciences, the University of Tokyo, which were originally collected from geographically and latitudinally different habitats and have been maintained for more than 30 years and many generations as closed colonies at an outdoor breeding facility (Shima, 1985a; Shima, 1985b). The parent fish used in this study were maintained as three independent pairs of 1 male and 1 female fish under standard laboratory conditions at 27.5 ± 1.5 °C with a 14:10 h light:dark photoperiod and were fed twice a day with live brine shrimp in the morning and commercial powder food (Otohime B1 and Tetrafin) in the evening.

### Low-temperature treatment and measurement of the heartbeat interval

Fertilized eggs were collected and incubated in plastic Petri dishes (diameter 60 mm) with tap water containing 0.0001% methylene blue for two days at 27 ± 0.2 °C. The designation of medaka developmental stages was in accordance with that of Iwamatsu (Iwamatsu, 1994). Embryos at st. 20, the four-somite stage, were transferred to an incubator at 15 ± 0.2 °C until st. 24, the stage at which heartbeat starts but without blood circulation. Embryos at st. 24 were used for the experiment.

Heartbeat measurement at st. 24 was performed once per embryo after incubation for 5 minutes at the random temperature in the range of 10–18 °C for Hd-rR strain and Kaga population and 12–15 °C for other medaka using a microscope (Multizoom AZ100; Nikon, Tokyo, Japan). Embryos were transferred to new incubation tap water containing 0.0001% methylene blue and data collection was performed without anesthetics because embryos at st.24 were in the chorion and enable observation under microscope. Heartbeat intervals were measured based on the time point of each heartbeat initiation. The onset of contraction of the heart was determined as the initiation time point of each heartbeat. The time intervals between adjacent heartbeats were measured for longer than one minute and more than ten heartbeats. The data collection was performed in one embryo for one temperature. The water temperature was measured with a handmade thermometer having a sensor (S-8100B; Seiko Epson Co., Nagano, Japan) calibrated against a standard thermometer. The water temperature was kept within 1 °C during the data collection for each embryo.

### Staging within the heartbeat initiation stage

Embryos of st. 24 were grouped into two stages based on the development of five tissues: heart, blood island, Kupffer’s vesicles, the otolith, and the blood cells in the blood vessels (Iwamatsu, 1994). The heart with a tubular shape or with atrium and ventricle, the presence or absence of a blood island, the presence or absence of Kupffer’s vesicles, the absence or presence of an otolith, and the absence or presence of blood cells in the blood vessels were categorized as “early” and “late,” respectively.

### Analyses based on averaged heart rate and CV

Extracted heartbeat intervals of each embryo were analyzed using two indices: heart rate and CV of interbeat interval, which was calculated as the standard deviation of the extracted heartbeat intervals divided by the averaged interbeat intervals. The mean value of heart rate and the CV of all the measured embryos in a strain were defined as heart rate and CV of the strain, respectively.

### Analyses based on temperature dependency

Heart rates and CVs were compared for each embryo in each strain according to the measurement temperature. Averaged heart rate and the heart rate variability of each medaka embryo within 12–15 °C were represented as a scatter plot of the average heart rate and CV by temperature, respectively. The heart rate and CV increment of individual embryos per 1 °C was obtained from an approximate straight line. Based on the slope of the approximate straight line from the scatter plot of measurement temperature and heart rate or CV, the temperature dependency of heart rate (dT_HR_ (beats/min/°C)) and of CV (dT_CV_ (/°C)) in each strain were evaluated for indicators of temperature dependence at low temperature.

### Statistical analyses

Statistical analyses were performed using Welch’s *t*-test and Tukey’s test. In this study, Welch’s *t*-test was used unless stated. A *P* value < 0.05 was considered to show significance. Data are presented as the mean ± standard deviation of the indicated samples per experiment.

## Results

### Temperature range of heartbeat analyses at low temperature

First, the heart rate and the CV of heartbeat intervals were examined for temperatures from 10 to 18 °C in embryos of the Kaga population derived from the N.JPN group and embryos of the Hd-rR strain derived from the S.JPN group. Regular heartbeat was observed in both medaka above 16 °C (Supplemental Figure 1A). All embryos in both strains showed similar constancy above 16 °C (Supplemental Figure 1B). The heartbeat stopped in most embryos of both medaka below 12 °C, suggesting that the cardiac pacemakers did not function in these embryos below 12 °C. Furthermore, we expected to find a threshold for the heart rate and its rhythm around 15 °C in the Hd-rR strain, so that cold tolerance of embryonic heart functions was different between the N.JPN and S.JPN groups. Therefore, each time interval between adjacent heartbeats was measured at 12–15 °C in the following experiments in this study.

### Comparison of embryonic heart rates at low temperatures among 13 local populations from the N.JPN, S.JPN, and W.KOR groups

The temperature and interbeat interval for each embryo were plotted (Figure 1) and average heart rates were calculated in 13 local populations of medaka (Figure 2). The heart rates of embryos of Kaga and Maizuru medaka in the N.JPN group were 8.8 ± 3.0 and 9.0 ± 3.1 beats/min (*P* > 0.05), respectively, and were higher than those of Odate embryos, which were 7.1 ± 2.6 beats/min (*P* < 0.05, Figures 1A – C and 2A). The interbeat interval and the average of heart rates of the S.JPN medaka were plotted in Figures 1D – K and 2B. The heart rate of Kazusa embryos was the highest in the S.JPN medaka at 12.3 ± 7.8 beats/min (*P* < 0.05). The heart rate of Arita and Urizura embryos was 9.2 ± 4.9 and 7.7 ± 4.4 beats/min (*P* > 0.05), which was higher than that of the embryos of the other S.JPN medaka (*P* < 0.05). The heart rate of Okaya embryos was 5.5 ± 1.8 beats/min (*P* < 0.05), and was as high as that of Ichinoseki, Iida, and Tanabe embryos, which were 6.4 ± 2.1, 6.4 ± 2.0, and 6.8 ± 2.3 beats/min, respectively (*P* > 0.05). There was no significant difference among the heart rates of Ichinoseki, Iida, and Tanabe embryos (*P* > 0.05), but were higher than that of Gushikami embryos, which was 4.5 ± 1.1 beats/min (*P* < 0.05).

**Figure 1.**
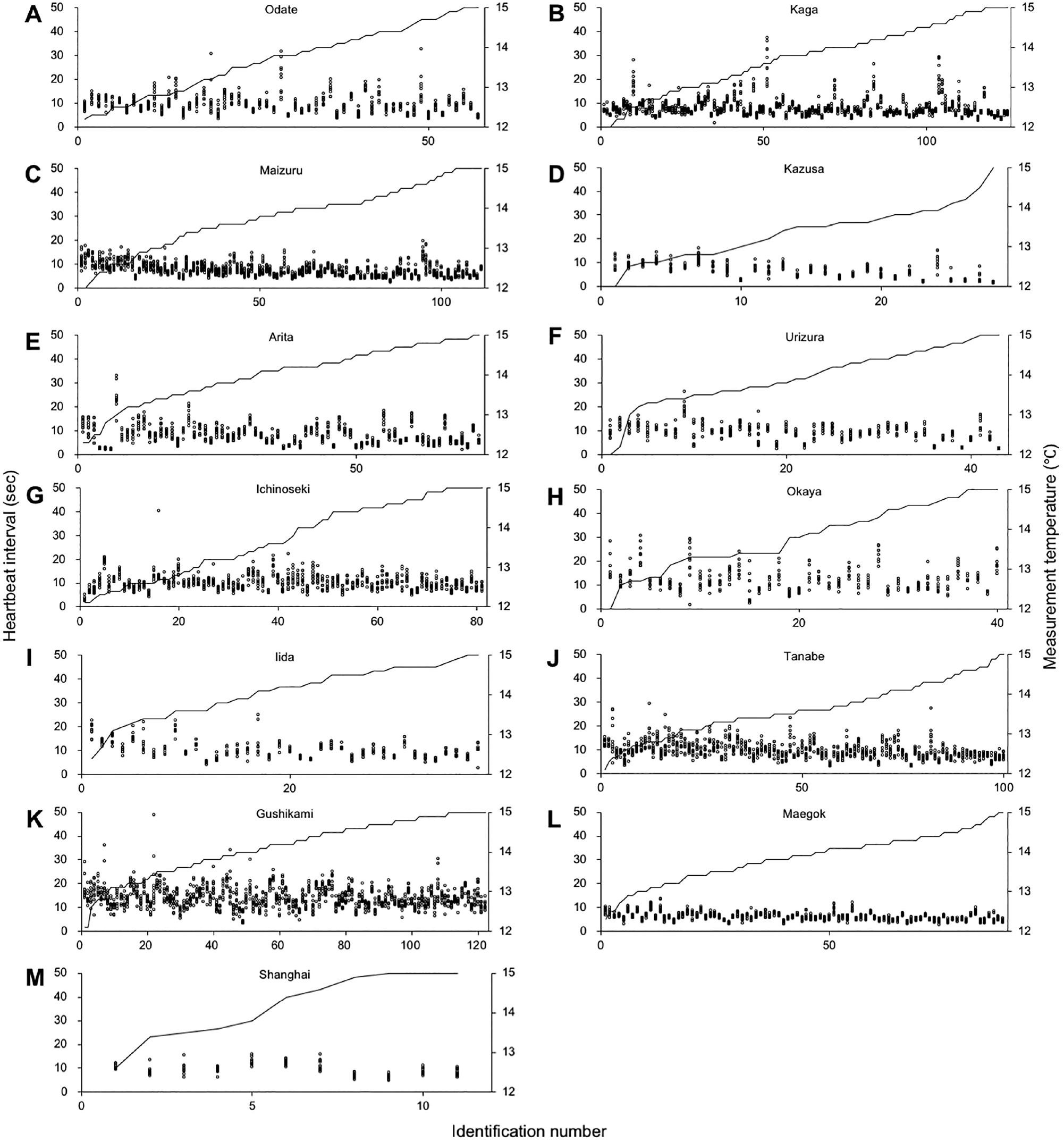
Heartbeat interval of medaka closed colonies under low temperature. The interbeat interval of embryonic medaka and its measurement temperature of each embryo in Odate (A), Kaga (B), Maizuru (C), Kazusa (D), Arita (E), Urizura (F), Ichinoseki (G), Okaya (H), Iida (I), Tanabe (J), Gushikami (K), Maegok (L), and Shanghai (M) medaka are shown in the initial ten interbeats for each embryo (dots in tandem) and in the line graph (secondary axis), respectively. Odate; n = 57, Kaga; n = 125, Maizuru; n = 111, Kazusa; n = 28, Arita; n = 72, Urizura; n = 43, Ichinoseki; n = 81, Okaya; n = 40, Iida; n = 38, Tanabe; n = 100, Gushikami; n = 122, Maegok; n = 88, Shanghai; n = 11.

**Figure 2.**
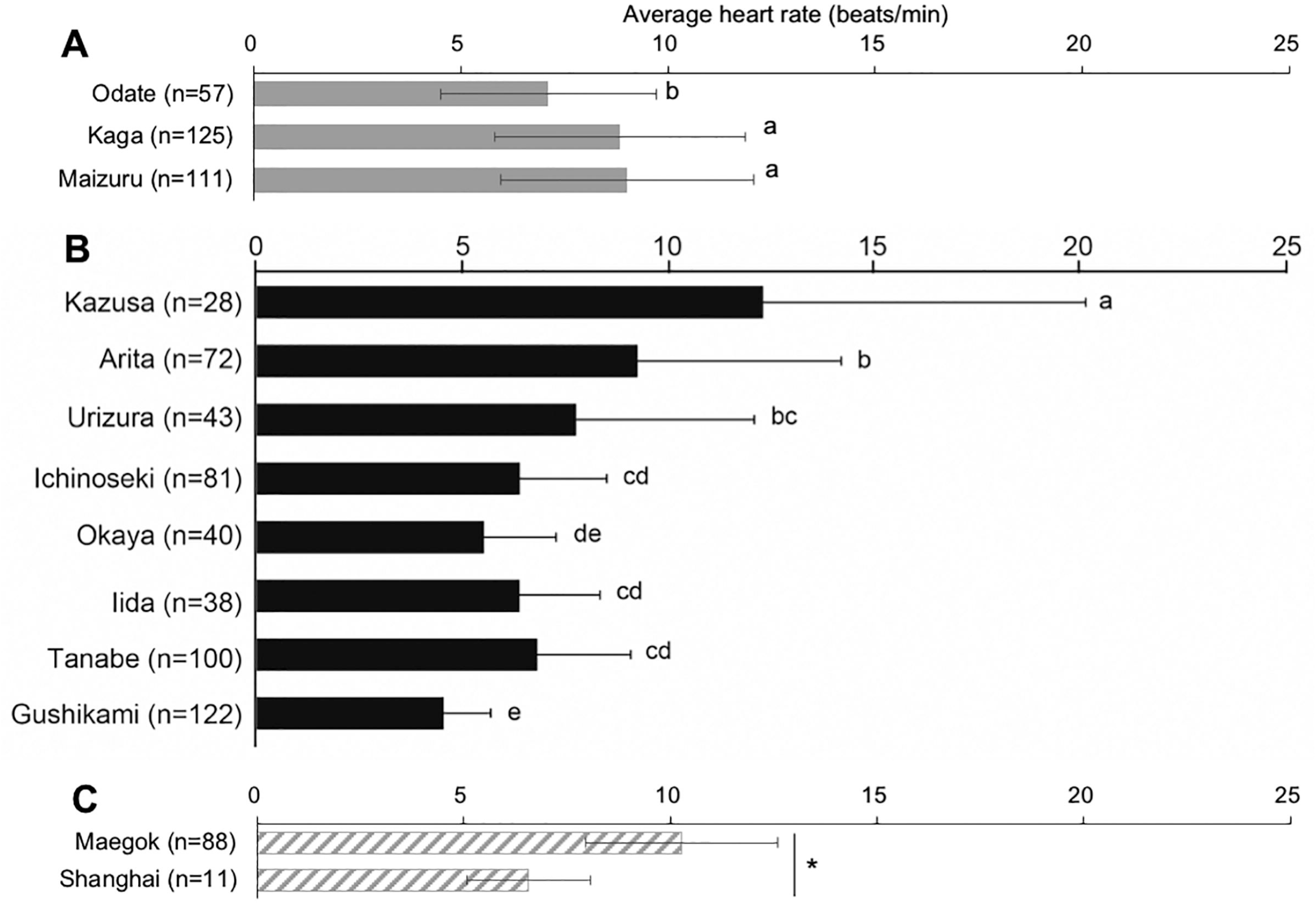
Comparison of heart rate at low temperatures in 13 wild populations. The heart rates of 13 medaka populations are compared in the N.JPN (A), S.JPN (B), and W.KOR groups (C). Bars labeled by the same letter (a, b, c, d, e) on the graph are not significantly different from each other (*P* > 0.05) by Tukey’s test. All data are presented as the means ± SD. (*: *P* < 0.01, n.s.: *P* > 0.05)..

The heart rate of embryos of Maegok medaka (the W.KOR group) was 10.3 ± 2.3 beats/min and was higher than that of Shanghai medaka embryos, which was 6.6 ± 1.5 beats/min (*P* < 0.0001, Figures 1L, M and 2C). These results demonstrate that heart rates varied among embryos from the same groups. Average heart rate of the three N.JPN, the eight S.JPN, and the two W.KOR medaka embryo groups were 8.3 ± 1.0, 7.3 ± 2.4, and 8.4 ± 2.6 beats/min, respectively, and there was no significant difference among them (*P* > 0.05, Supplemental Figure 2A). These results suggest that average heart rates of st. 24 embryos at low temperatures were not determined genetically by the group but varied among the populations.

The heart rate varies depending on the stage of embryonic development, especially around the heartbeat initiation stage (st. 24) at 25.5 °C (Matsui, 1941). We compared the heart rate according to the developmental stage of embryos in st. 24 (supplemental Figure 3A). The average heart rate of late st. 24 embryos was higher than that of early st. 24 embryos in Kaga, Odate, Tanabe (*P* < 0.05), and Maegok medaka (*P* < 0.01), but showed no significant difference from that of early st. 24 in Maizuru (*P* > 0.05). These results suggest that the heart rate varied at the developmental stage according to the medaka populations, while there was a tendency for the heart rate to increase with the embryonic development in all wild populations examined. There was no significant difference in the average heart rate in st. 24 embryos with development among the N.JPN, S.JPN, and W.KOR groups.

### Comparison of the CV at low temperatures in 13 wild populations from the N.JPN, S.JPN, and W.KOR groups

We next compared the variability of the heartbeat interval among the embryos of local populations by comparing the CV of interbeat intervals in st. 24 embryos. In the N.JPN group, the CV of Odate and Maizuru embryos were 0.17 ± 0.09 and 0.16 ± 0.07 (*P* > 0.05, Figure 3A), respectively, and heartbeat intervals were more consistent in both populations than in Kaga embryos, for which the CV was 0.13 ± 0.06 (*P* < 0.05). In the S.JPN medaka, the CV of Okaya and Tanabe embryos were 0.21 ± 0.12 and 0.19 ± 0.09 (*P* > 0.05), respectively, higher than that of Iida medaka, which was 0.14 ± 0.08 (*P* < 0.05, Figure 3B). The CV of Ichinoseki, Urizura, Arita, Kazusa, and Gushikami embryos were 0.17 ± 0.07, 0.16 ± 0.11, 0.16 ± 0.07, 0.18 ± 0.10, and 0.17 ± 0.07, respectively, but there was no significant difference among them (*P* > 0.05, Figure 3B). The CV of Maegok medaka in the W.KOR was 0.10 ± 0.04, lower than that of Shanghai medaka, which was 0.16 ± 0.05 (*P* < 0.005, Figure 3C). These results strongly suggest that the CV within 12–15 °C was variable among local populations and there was no tendency among the genetically distinctive three groups. The mean value of the CV of three N.JPN, eight S.JPN, and two W.KOR medaka were 0.15 ± 0.02, 0.17 ± 0.02, and 0.13 ± 0.04, respectively, and there was no significant difference among them (*P* > 0.05, Supplemental Figure 2B). These results confirm that the mean value of the CV at st. 24 within 12–15 °C was not different in the three groups.

**Figure. 3.**
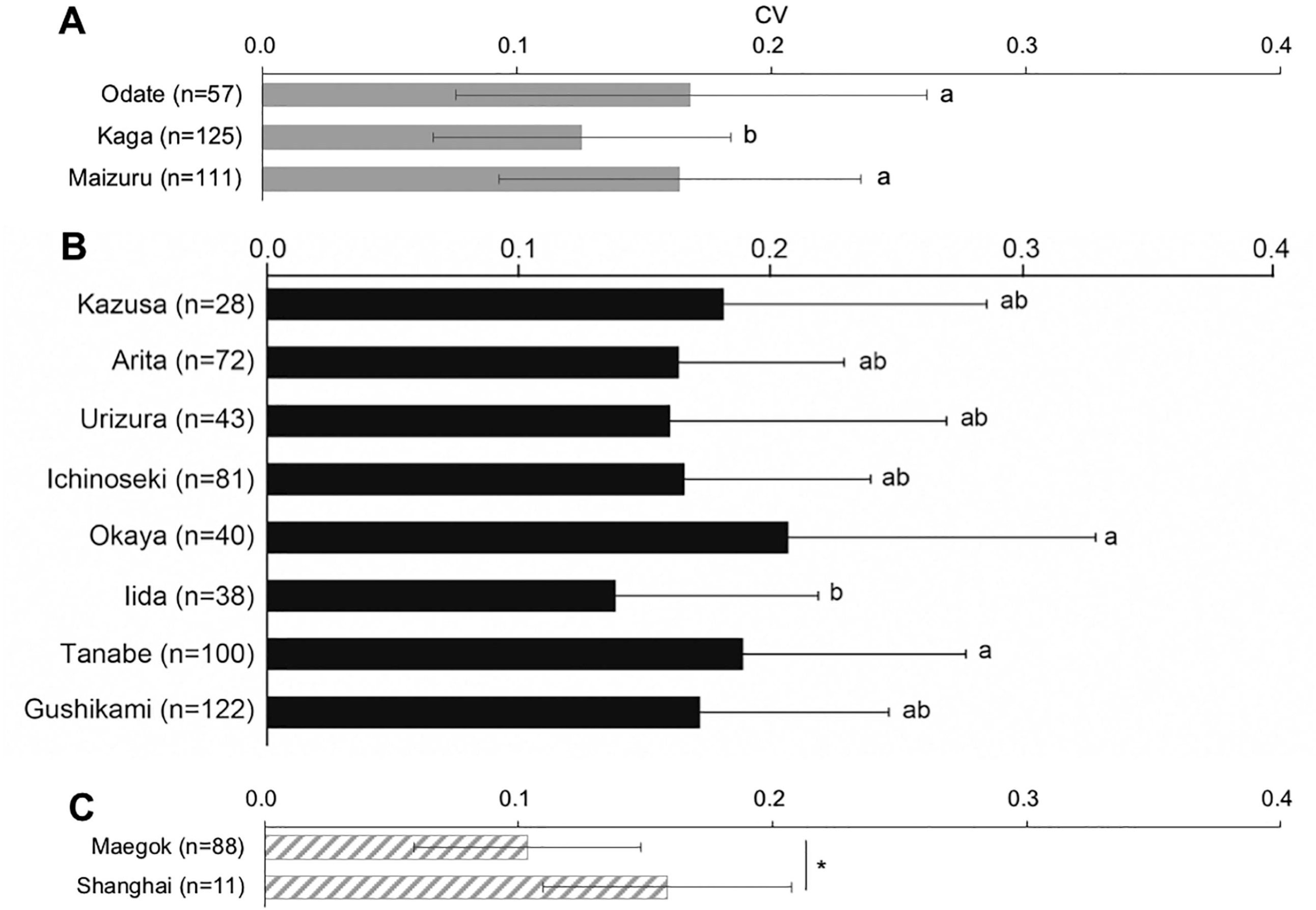
Comparison of the CV at low temperatures in 13 wild populations. The CV of medaka embryos was compared in the N.JPN (A), S.JPN (B), and W.KOR groups (C). Bars labeled by the same letter (a, b) on the graphs are not significantly different from each other (*P* > 0.05) by Tukey’s test. All data are presented as the means ± SD. (*: *P* < 0.01, n.s.: *P* > 0.05). All data are presented as the means ± SD. (*: *P* < 0.01, n.s.: *P* > 0.05).

We next compared the variation in heart rate using the standard deviation of heartbeat interval (SD_HR_) according to the developmental stage of embryo in st. 24 (supplemental Fig. 3B) and found that there was no significant difference between early and late st. 24 in most of the medaka populations (*P* > 0.05), suggesting that the SD_HR_ was irrelevant to the developmental stage.

### The temperature dependency of heart rate and CV at low temperatures

We next analyzed the temperature dependency of embryonic heart physiology within 12–15 °C (Figure 4). The heart rate decreasement of embryos per 1 °C within 12–15 °C (dT_HR_) was calculated as the temperature dependency of heart rate (Figure 4A). The dT_HR_ of the Odate, Kaga, and Maizuru embryos in the N.JPN group were 0.77, 0.75, and 2.01 beats/min/°C, respectively. The dT_HR_ of the Ichinoseki, Urizura, Okaya, Iida, Tanabe, Arita, Kazusa, and Gushikami embryos in the S.JPN group were –0.15, 2.46, 0.39, 1.17, 1.49, 0.44, 8.56, and 0.25 beats/min/°C, respectively. The dT_HR_ of Maegok and Shanghai embryos were 1.57 and 0.84 beats/min/°C, respectively. These results suggested that there was no tendency in the temperature dependence except Kazusa population.

**Figure. 4.**
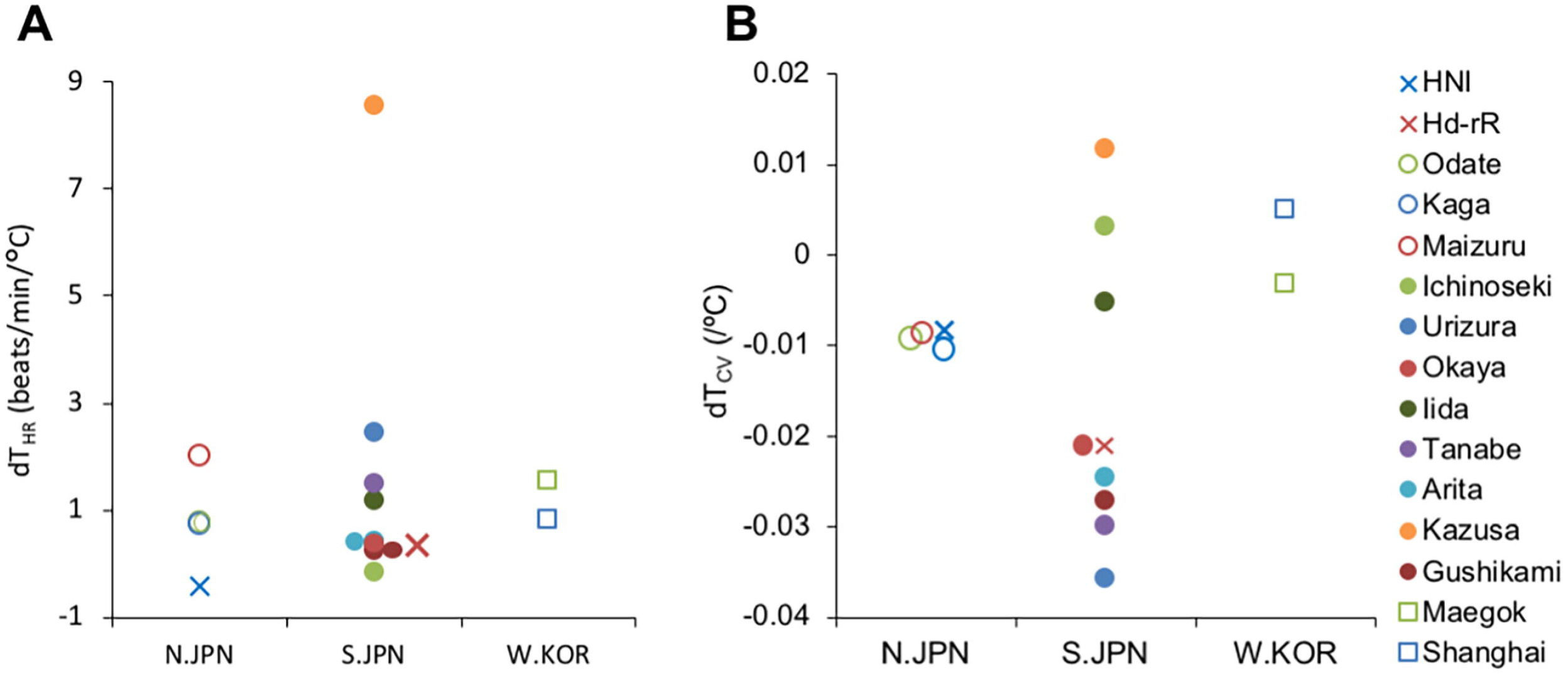
Comparison of the average heart rate and the CV in the three groups and temperature dependency at low temperatures in 13 wild populations. The heart rate (A) and the CV (C) of 13 wild populations were averaged in its respective group; N.JPN, S.JPN or W.KOR. The slopes of the approximate straight lines from the scatter plots of measurement temperature and the heart rate (dT_HR_) and the CV (dT_CV_) are compared as a temperature dependency of the heart rate (B) and the CV (D) in the N.JPN, S.JPN, and W.KOR groups including the inbred strains, respectively. All data are presented as the means ± SD. (*: *P* < 0.01, n.s.: *P* > 0.05). The population of N.JPN, S.JPN, W.KOR and inbred strains indicated in open circles, solid circles, squares and crosses, respectively. HNI, n = 46; Hd-rR, n = 48; Odate, n = 57; Kaga, n = 125; Maizuru, n = 111; Ichinoseki, n = 81; Urizura, n = 43; Okaya, n = 40; Iida, n = 38; Tanabe, n = 100; Arita, n = 72; Kazusa, n = 28; Gushikami, n = 122; Maegok, n = 88; Shanghai, n = 11.

The CV increment of embryos per 1 °C within 12–15 °C (dT_CV_) was calculated as the temperature dependency of CV (Figure 4B). The dT_CV_ of the Odate, Kaga, and Maizuru embryos in the N.JPN medaka were –0.008, –0.010, and –0.008/°C, respectively. The dT_CV_ of the Ichinoseki, Urizura, Okaya, Iida, Tanabe, Arita, Kazusa, and Gushikami embryos in the S.JPN group were 0.003, –0.036, –0.021, –0.005, –0.030, –0.025, 0.012, and –0.027/°C, respectively. The dT_CV_ of Maegok and Shanghai in the W.KOR group were –0.003 and 0.005/°C, respectively. In the S.JPN group, Ichinoseki, Iida, and Kazusa embryos showed higher dT_CV_ values than the embryos of the other S.JPN medaka. These results suggested that the dT_CV_ values of the S.JPN groups showed existence of CV variation in temperature dependency only in the S.JPN embryos.

### Heartbeat at low temperature of two inbred strains from the N.JPN and S.JPN groups

A series of ten heartbeats (open circles) in one embryo was listed vertically for each embryo and arranged in ascending order of measurement temperature within 12–15 °C (Figures 5A and B). Averages of heart rates of HNI and Hd-rR embryos were 5.6 ± 2.5 and 3.0 ± 0.9 beats/min (*P* < 0.0001, Figure 5C), and the CVs of HNI and Hd-rR embryos were 0.13 ± 0.05 and 0.36 ± 0.11 (*P* < 0.0001, Figure 5D), respectively. The dT_HR_ of the of Hd-rR and HNI strain was 0.38 and –0.42 beats/min/°C, respectively (Figures 4A, 5E and F). These results confirmed the results from the wild populations. The dT_CV_ of the Hd-rR and the HNI strain were –0.021 and –0.008/°C, respectively (Figures 4B, 5G and H). These results suggest that temperature dependencies of the heart rate and its CV were the phenotype characteristic to the S.JPN medaka embryo and were undeniable to results of wild populations.

**Figure 5.**
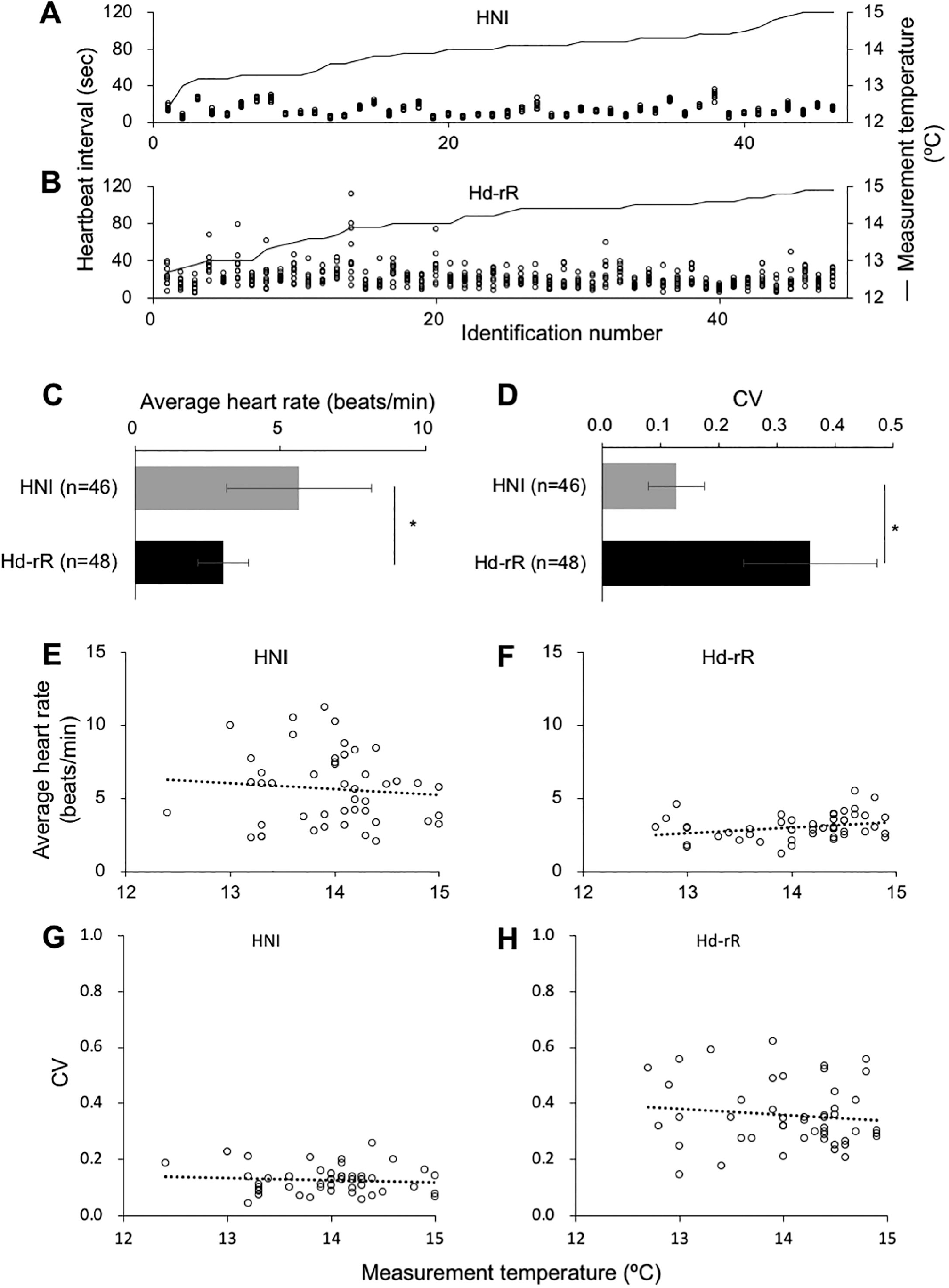
Comparison of heartbeat at low temperatures between two inbred strains. Heartbeat interval of embryonic medaka and the measurement temperature of each embryo in HNI (A) and Hd-rR (B) strains are shown in the initial ten interbeats for each embryo (dots in tandem) and in the line graph (secondary axis), respectively. The heart rate (C) and the CV (D) are compared between the two strains. The heart rate and the CV of each embryo are compared for each measurement temperature and shown in a scatter plot of measurement temperature and heart rate or CV in HNI (E, G) and Hd-rR (F, H) strains. All data are presented as means ± SD. (*: *P* < 0.01). HNI, n = 46; Hd-rR, n = 48.

## Discussion

These results suggest that average heart rates of st. 24 embryos at low temperatures were not determined genetically by the group but varied among the populations. Therefore, the difference between the heart rates of the HNI and Hd-rR embryos shown in Figure 2C were the characteristics of HNI and Hd-rR embryos and did not represent those of the N.JPN and S.JPN medaka groups.

It is noteworthy that the heart rate above 17 °C was temperature dependent in all medaka populations (Matsui, 1941). The N.JPN, S.JPN, and W.KOR groups are genetically different, consisting of distinct local populations (Takehana, 2003). Our results show no strong temperature tendency or specific differences in the cold tolerance of the embryonic heart physiology among the N.JPN and W.KOR group medaka; however, the S.JPN showed two different phenotypes: Kazusa medaka with a strong temperature dependency, and others without strong temperature dependency at 12–15 °C. We consider that the positive correlation in dT_HR_ of Kazusa embryos was caused by the rapid heartbeat around 15 °C (supplemental Figure 1J).

These results clearly demonstrate that the two strains and the 13 local populations of medaka can be divided into two groups: a “negative-correlation group,” which includes the Hd-rR strain, Urizura, Okaya, Tanabe, Arita, and Gushikami medaka in the S.JPN group and a “less-correlation group,” which includes the HNI strain, Odate, Kaga, and Maizuru medaka in the N.JPN group, Ichinoseki, Iida, and Kazusa medaka in the S.JPN group, and Maegok and Shanghai medaka in the W.KOR group by the evaluation of temperature dependency of the CV within the range of 12–15 °C. These findings also suggest that the N.JPN, S.JPN, and W.KOR medaka groups showed specific differences in the temperature dependency of the CV of heartbeat interval in st. 24 embryos, while there was no difference in the average heart rate, the CV of heartbeat interval, nor any temperature dependency of heart rate (dT_HR_) in temperature range of 12–15 °C.

The heart rate of Odate (N.JPN) embryos was more stable than that of Hd-rR (S.JPN) embryos at st. 24 at 16 °C (Watanabe-Asaka, 2014). In the previous study, both S.JPN medaka and W.KOR Shanghai medaka showed blood regurgitation at st. 36, but the N.JPN medaka did not, suggesting that the N.JPN acquired a stable heartbeat during embryogenesis at low temperature. This might allow the N.JPN to expand to the northern part of Japan. On the other hand, this study demonstrated that the N.JPN and W.KOR groups, including Shanghai medaka, showed stable heart rates at low temperatures and the S.JPN group showed both stable and increased heart rate variability with decreasing temperature. These contradictory results in the W.KOR medaka indicated that the blood regurgitation at st. 36 was caused not only by the bradyarrhythmia at st. 24 but also by other dysfunctions during embryogenesis before st. 36 such as retardation of angiogenesis or impairment of hemodynamic forces (Andrés-Delgado, 2016; Collins, 2016).

We consider that the difference in heart rate variability at low temperatures among the populations was caused by the different genetic backgrounds of medaka rather than by environmental effects, because all medaka populations examined in this study have been maintained for generations as closed colonies at the same outdoor breeding facility. It was widely accepted that the two regional Japanese populations, the N.JPN and S.JPN groups, have diverged after the W.KOR group from the common ancestor of these three groups (Watanabe, 2006; Kasahara, 2007). We may therefore propose that the ancestral population of *O. latipes*, which had cold tolerance, spread to higher latitudes in East Asia from the low-latitude areas, and the ancestral *O. latipes* population might have diverged.

According to the neighbor-joining tree of the entire cytochrome *b* gene in the S.JPN group (Takehana, 2003), Kazusa, Ichinoseki, and Iida medaka, which had CVs that were not temperature dependent like the N.JPN and W.KOR medaka, were classified into the subclades B-XI, B-I and B-I, respectively. However, the Hd-rR strain and the Urizura, Okaya, Tanabe, Arita, and Gushikami medaka, which had temperature-dependent CVs, were classified into the subclades B-II, B-I, B-I, B-VII, B-XI and B-XI, respectively. When we consider the sharing of the temperature dependency of CV among the S.JPN medaka in the different latitudes and with the different genetic backgrounds, the following scenario of natural history of medaka in the Japan archipelago can be proposed. The ancestral medaka population, which is common to the four wild medaka groups (W.KOR, E.KOR, N.JPN, and S.JPN) expanded to high latitudes in mainland China with their cold tolerance and reached the Korean Peninsula (Watanabe, 2006). Then, part of the population further expanded to the Japan archipelago and divided into the two groups that were the direct ancestors of the N.JPN and S.JPN medaka with their cold tolerance. While the N.JPN medaka inhabited the cold-climate area and have retained their cold tolerance up to the present era, some populations of the S.JPN group lost their cold tolerance because of the loss of selective pressure in their warm-climate habitats and kept varying until the present era. It is widely thought that the N.JPN medaka acquired their cold tolerance and expanded to the cold climate habitats in the Japan archipelago; however, the findings reported here suggest that cold tolerance is an intrinsic feature of medaka and some of the S.JPN medaka lost it rather than that the medaka populations of the S.JPN group acquired their cold tolerance independently.

## Conclusions

In this study, we have analyzed the heart rates of embryos of three N.JPN, eight S.JPN, and two W.KOR medaka closed colonies at low temperatures (within 12–15 °C). Our findings suggest that the N.JPN, S.JPN, and W.KOR medaka groups showed specific differences in the temperature dependency of the CV of heartbeat interval in st. 24 embryos, while there was no difference in the average heart rate, the CV of heartbeat intervals, nor the temperature dependency of heart rate (dT_HR_) in this temperature range. The difference in heart rate variability at low temperatures among the wild populations was caused by the different genetic background of medaka rather than environmental effects. The findings in this study confirm that cold tolerance is an intrinsic feature of medaka, but that some of the S.JPN medaka lost it. These findings provide cues for the natural history of the S.JPN medaka inhabiting the Japanese archipelago.

## Declarations

### Ethics approval and consent to participate

Not applicable

### Availability of data and materials

The datasets used and/or analyzed during the current study are available from the corresponding author on reasonable request.

### Authors’ contributions

H. M. and T. W.-A. designed the experiments; W. Y., S. O., and T.W.-A. performed the experiments; W.Y., S.O., T.K., and T. W.-A. analysed the data; S. O., T. K., H. M. and T. W.-A. interpreted the results of the experiments; S. O. and T. W.-A. drafted the manuscript; W. Y., S. O., T. K., H. M., and T.W.-A. edited the manuscript. S. O., H. M., and T.W.-A. revised the manuscript. All authors have read and approved the final manuscript.

## Acknowledgements

We express our appreciation to Mmes. Shizuko Chiba, Sumiko Tomizuka, Ikumi Matsumoto and Mayuko Takagi for maintaining of medaka strains and closed colonies. The authors thank NBRP for providing the wild medaka lab stocks.

