## Supplementary figures and images for "Intraspecific diversity based on low temperature induced arrhythmia in embryonic heart of medaka (*Oryzias latipes*)"

### Supplemental Fig. 1

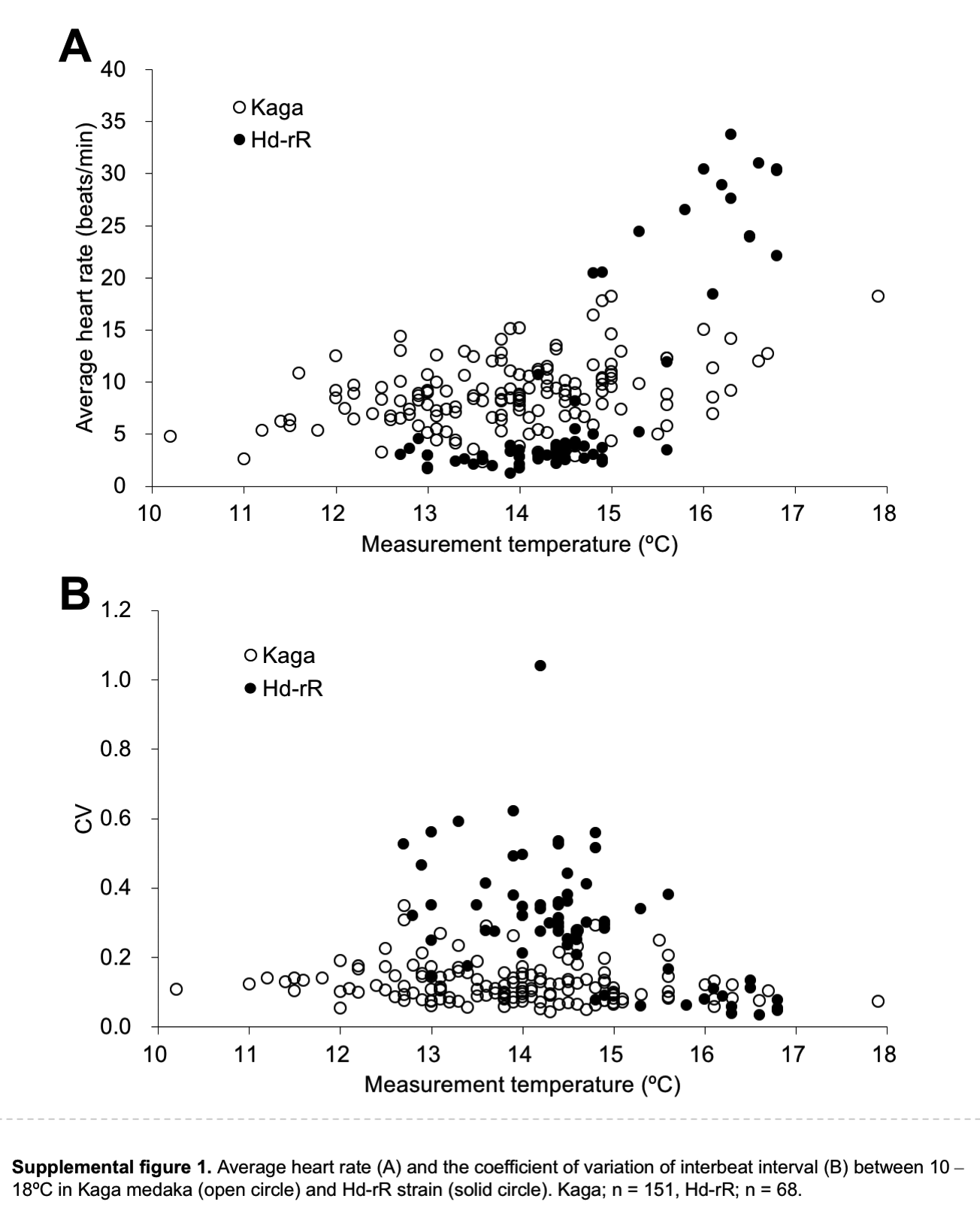

### Supplemental Fig. 2

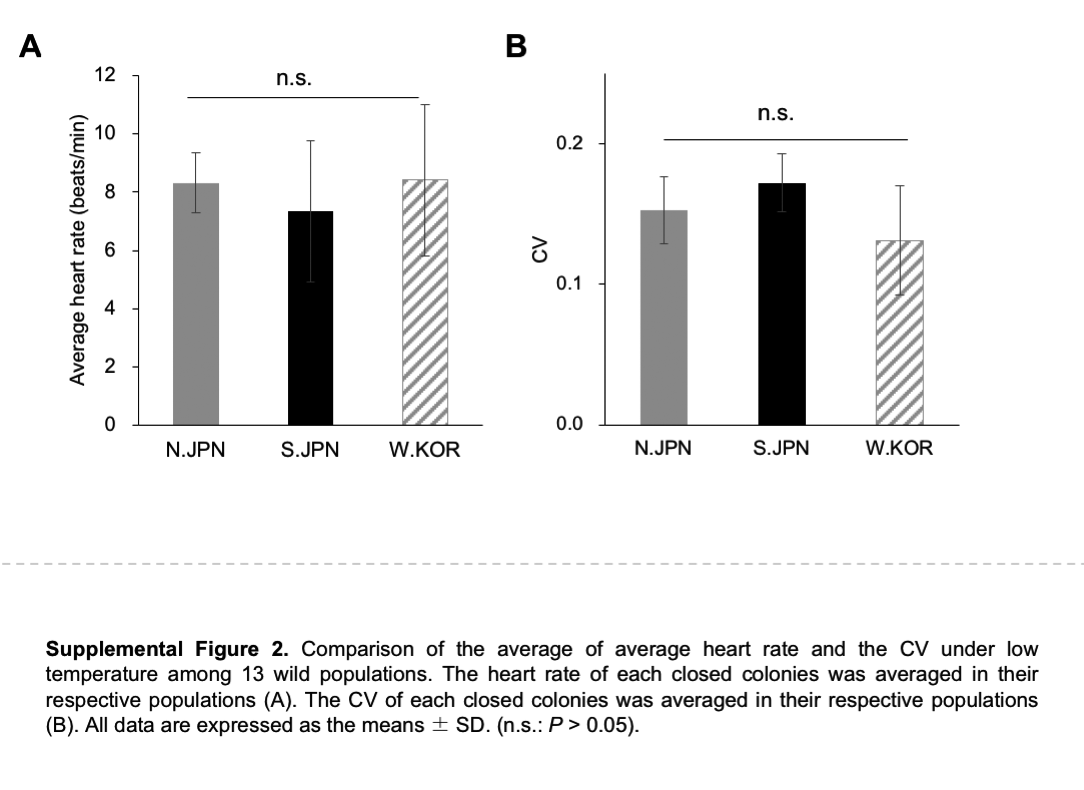

### Supplemental Fig. 3

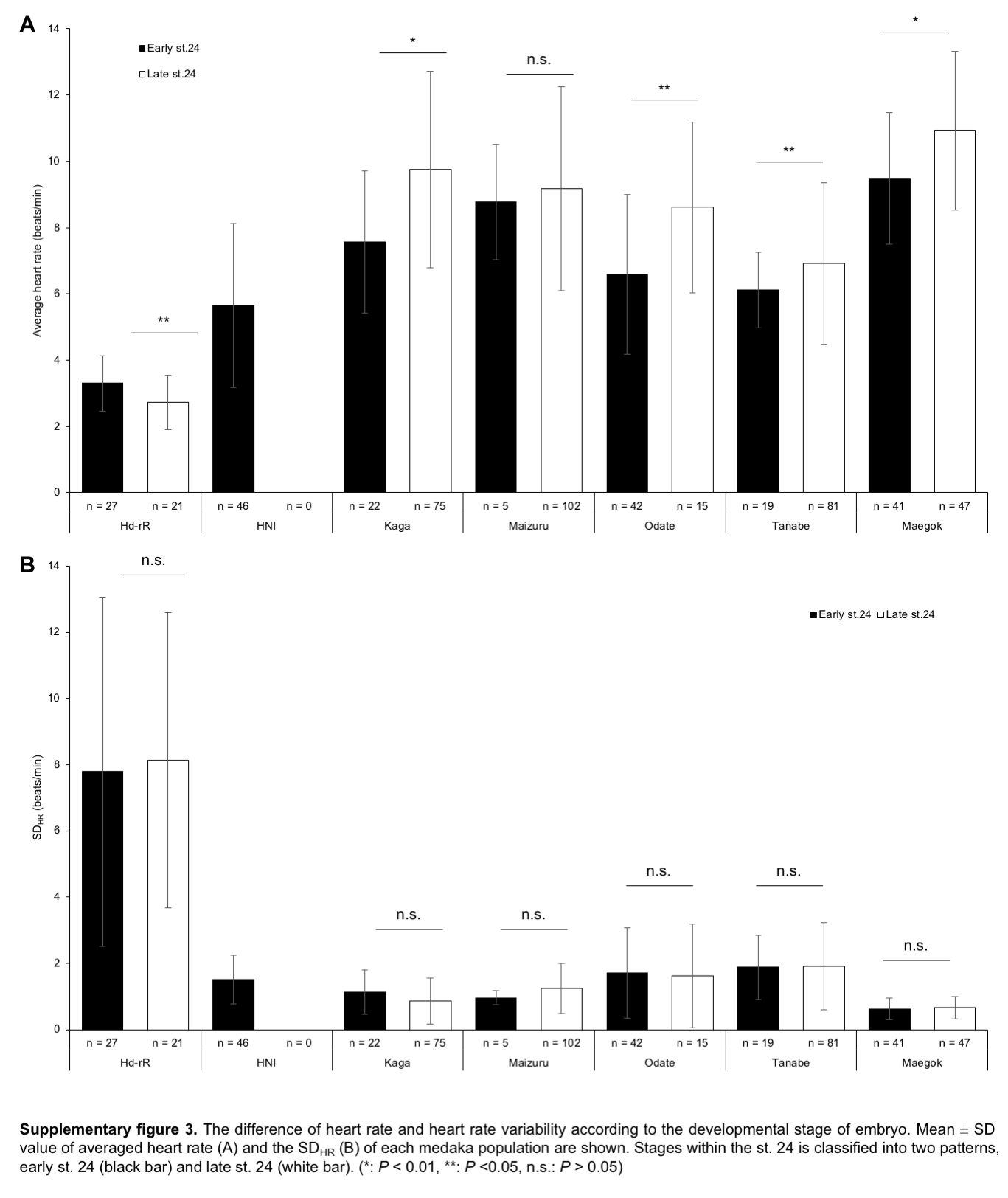
